# Genome report: A duplication lost in sugarcane hybrids revealed by chloroplast genome assembly of wild species *Saccharum officinarum*

**DOI:** 10.1101/141002

**Authors:** Deise Paes, Filipe Pereira Matteoli, Thiago Motta Venancio, Paulo Cavalcanti Gomes Ferreira, Clicia Grativol

**Author notes:** **Corresponding author**: Clicia Grativol Laboratório de Química e Função de Proteínas e Peptídeos, Centro de Biociências e Biotecnologia, Universidade Estadual do Norte Fluminense Darcy Ribeiro, Av. Alberto Lamego, 2000, P5-228, Parque Califórnia, Campos dos Goytacazes-RJ, 28013-602, Brazil.

## Abstract

Sugarcane is a crop of paramount importance for sustainable energy. Modern sugarcane cultivars are derived from interspecific crosses between the two wild species *Saccharum officinarum* and *Saccharum spontaneum* and this event occurred very early in the sugarcane domestication history. This hybridization allowed the generation of cultivars with complex aneuploidy genomes containing 100–130 chromosomes that are unequally inherited - ~80% from *S. officinarum*, ~10% from *S. spontaneum* and ~10% from inter-specific crosses. Several studies have highlighted the importance of chloroplast genomes (cpDNA) to investigate hybridization events in plant lineages. Few sugarcane cpDNAs have been assembled and published, including those from sugarcane hybrids. However, cpDNAs of wild Saccharum species remains unexplored. In the present study, we used whole-genome sequencing data to survey the chloroplast genome of the wild sugarcane species *S. officinarum*. Illumina sequencing technology was used for assembly 142,234 bp of *S.officinarum* cpDNA with 2,065,893 reads and 1043x of coverage. The analysis of the *S. officinarum* cpDNA revealed a notable difference in the LSC region of wild and cultivated sugarcanes. Chloroplasts of sugarcane cultivars showed a loss of a duplicated fragment with 1,031 bp in the beginning of the LSC region, which decreased the chloroplast gene content in hybrids. Based on these results, we propose the comparative analysis of organelle genomes as a very important tool for deciphering and understanding hybrid Saccharum lineages.

## INTRODUCTION

Sequencing of organelle genomes is an important tool in molecular and evolutionary studies (Wolf *et al*. 2011). In addition to the nuclear genome, plants have mitochondrial (mtDNA) and chloroplast genomes (cpDNA), which can allow a broad analysis on specific species (Xu *et al*. 2015). The size of cpDNAs of land plants ranges from 100 to 160 kb, with around 100 to 120 highly conserved genes (Wicke *et al*. 2011; Olejniczak *et al*. 2016). The features of the cpDNAs are also helpful in phylogenetic studies and to develop genetic markers (Bock and Khan 2004; Jansen *et al*. 2007; Ravi *et al*. 2008; Wu and Ge 2012). Further, several studies highlighted the importance of cpDNAs to investigate hybridization events than nuclear genomes; cpDNAs allow the analysis of organelle sharing patterns between species due to their slow rate of evolution, non-recombinant nature, easy haplotype detection and predominantly uniparental inheritance (Wu *et al*. 2010; Smith 2015; Zhu *et al*. 2016; Szczecińska *et al*. 2017; Xiao-Ming *et al*. 2017). Many studies have been conducted with chloroplast genomes to identify the history of plant lineages (Marí-ordóñez *et al*. 2013; Rousseau-Gueutin *et al*. 2015; Cho *et al*. 2016; Shetty *et al*. 2016; Yang *et al*. 2016; Asaf *et al*. 2017). As an example, the sequencing of chloroplast genomes of *Solanum commersonii* and *Solanum tuberosum* revealed indel markers that can distinguish chlorotypes and maternal inheritance of these organelles in hybrids (Cho *et al*. 2016).

Many species from the *Saccharum* genus (Poaceae) have been widely used in sugar production due to their remarkable sucrose storage capacity. Due to its tropical and subtropical distribution, sugarcane has probably been first established at New Guinea and Indonesia (Grivet *et al*. 2006). *S. officinarum* has a chromosome number of 2n = 80 and is known as “noble” sugarcane, mainly due to its high sucrose content, large and thick low-fiber stalks (Cheavegatti-Gianotto et al., 2011). Despite these key agronomic traits, this species is water-intensive, susceptible to diseases and requires high soil fertility. In the end of 19 century, a cross between *Saccharum spontaneum* and *Saccharum officinarum* resulted in a hybrid that was then backcrossed with *Saccharum officinarum*. The introgression of a small part of the *S. spontaneum* genome into a predominantly *S. officinarum* genome resulted in modern hybrids *(Saccharum spp.)* with better yields, high sucrose content and ability to cope some biotic and abiotic stresses. These hybrids were critical for the development of the sugar trade (Grivet and Arruda 2002; Moore 2005; Cheavegatti-Gianotto *et al*. 2011). Modern sugarcane cultivars have complex and aneuploidy nuclear genomes. Few sugarcane cpDNAs have been assembled and published, including those from the sugarcane hybrids *Saccharum* spp. Q155 (Hoang *et al*. 2015), *Saccharum* spp. NCo 310 (Asano *et al*. 2004), *Saccharum* spp. SP80-3280 (Calsa Júnior *et al*. 2004) and *Saccharum* spp. RB867515 (Vidigal *et al*. 2016). However, cpDNAs of wild Saccharum species remains unexplored. In the present study, we used whole-genome sequencing data to survey the chloroplast genome of the wild sugarcane species *S. officinarum*.

## METHODS AND MATERIALS

### Plant material

Young leaves from *S. officinarum* accession 82-72 maintained in the germplasm collection of Instituto Agronômico de Campinas (Ribeirão Preto, Brazil) were used for DNA analysis. According to Kuijper’s leaf numbering system for sugarcane (Cheavegatti-Gianotto *et al*. 2011), leaf -2 tissue was used to subsequent DNA extraction and sequencing.

### DNA extraction and sequencing

Total genomic DNA was extracted from leaves using the CTAB method (Doyle and Doyle 1987) with minor modifications. The quality of DNA was estimated using Thermo Scientific *NanoDrop*^™^ 2000c Spectrophotometer. Total DNA (~20ug) was sequenced on the Illumina GAII machine using the paired-end 100 cycle protocol.

### *De novo* assembly of chloroplast using genomic DNA reads

The sequencing reads were initially filtered to retain those with 90% of bases having quality scores greater than or equal to 20 (Q20) using FASTX Toolkit (http://hannonlab.cshl.edu/fastx_toolkit/). After quality filtering, we performed a BLASTN (Altschul *et al*. 1997) search (e-value ≤ 10^−4^) with chloroplast sequences from *Saccharum* hybrid cultivar NCo 310 (NC_006084.1), *Saccharum* hybrid cultivar SP-80-3280 (AE009947.2), *Sorghum bicolor* (NC_008602.1), *Zea mays* (NC_001666.2), *Miscanthus sinensis* (NC_028721.1), *Oryza sativa* (KT289404.1) and *Setaria italica* (KJ001642.1). The reads aligned to cpDNAs were tested on VelvetOptimiser (https://github.com/tseemann/VelvetOptimiser) with k-mer range from 29 to 87. Genome assembly was performed with SPADES (Bankevich *et al*. 2012) using the following parameters: 53, 69 and 77 of k-mers; 70 of coverage cutoff and careful parameter. The SSPACE (Boetzer *et al*. 2011) was run with default parameters on SPADES assembled contigs. The assembled chloroplast was compared with cpDNA from Saccharum hybrid cultivar Q155 using BLASTN, annotated with GeSeq (Tillich *et al*. 2017) and the resulting Genbank file was visualized on OrganellarGenomeDRAW (Lohse *et al*. 2013).

### Data availability

The sequence data from whole genome shotgun of sugarcane wild species *S. officinarum* have been submitted to the NCBI Sequence Read Archive under accession SRX313496. The assembled chloroplast genome sequence is available at NCBI Genbank with accession number MF140336 (http://www.ncbi.nlm.nih.gov/nuccore/MF140336).

## RESULTS AND DISCUSSION

A total 297,637,906 whole-genome shotgun reads of *S. officinarum* were sequenced using an Illumina GAII platform. Quality filtered reads were screened for similarity with known chloroplast sequences (see methods for details), which resulted in 2,065,893 reads that were used to assemble the 142,234 bp *S. officinarum* cpDNA at 1043x coverage (Table 1). Five scaffolds were assembled, the largest one with 106,869 bp (Table 1). This genome has 1,052 bp more than the hybrid cultivar *Saccharum spp*. SP80-3280, reported to have 141,182 bp (Calsa Júnior *et al*. 2004). The *S. officinarum* cpDNA has four main regions: LSC and SSC, with 84,080 bp and 12,576 bp, respectively and; the inverted repeats, IRa and IRb, with 22,789 bp each (Figure 1). Seventy-two genes were annotated, out of which 25 are protein-coding genes, 40 tRNA genes, four rRNA and three other genes (cssa, cemA and infA). In the inverted region, there are 20 duplicate genes: ten tRNA, four rRNA and six protein-coding genes. The IR junction with LSC is between the *rp122* and *trnH-rps19* gene cluster. Accordingly, the *trnH-rps19* gene cluster is present close to in the IR/LSC junction region in other monocotyledons species chloroplasts (Wang *et al*. 2008).

**Table 1.**
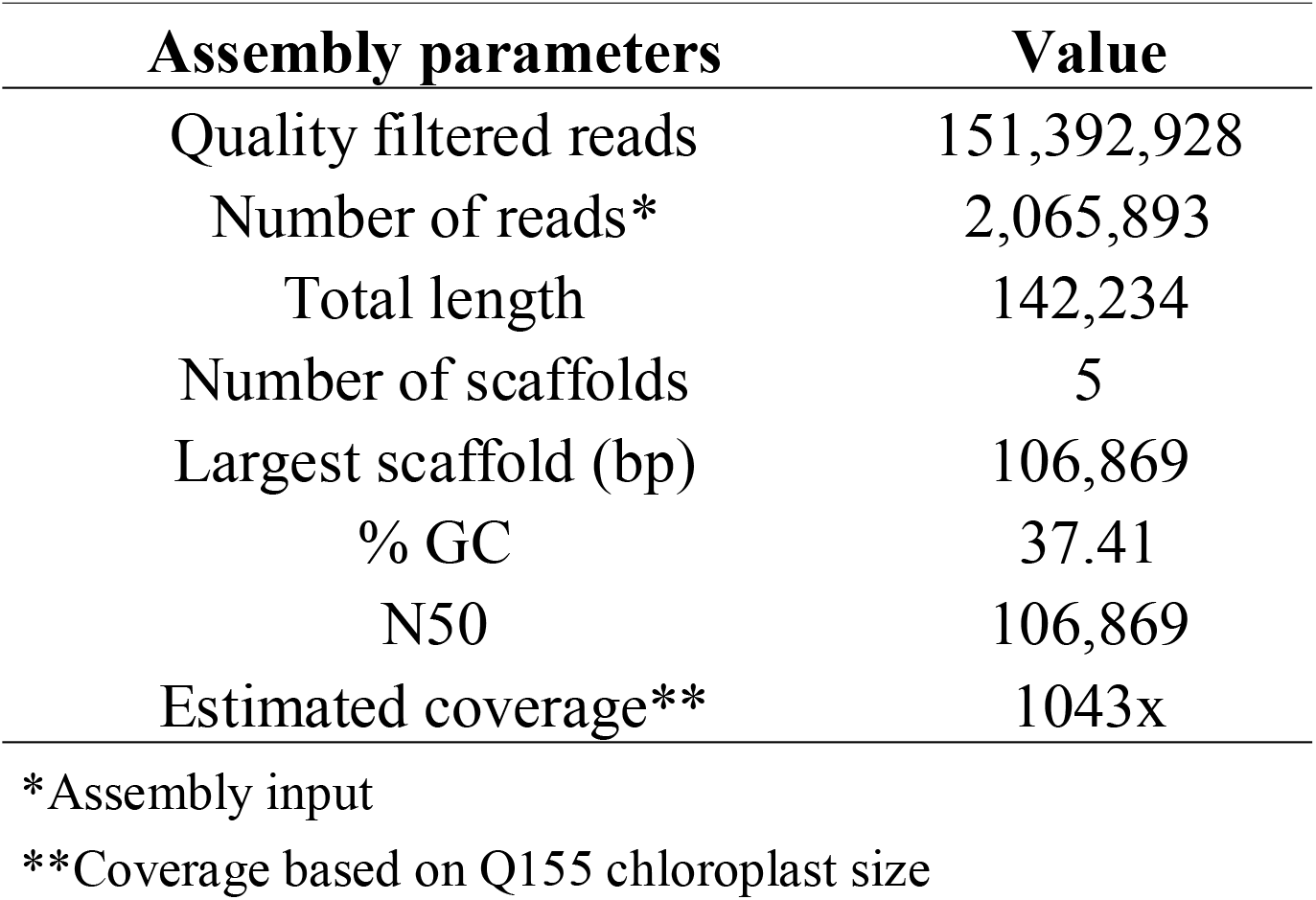
Summary of *S. officinarum* chloroplast genome assembly statistics.

| **Assembly parameters** | **Value** |
| --- | --- |
| Quality filtered reads | 151,392,928 |
| Number of reads* | 2,065,893 |
| Total length | 142,234 |
| Number of scaffolds | 5 |
| Largest scaffold (bp) | 106,869 |
| % GC | 37.41 |
| N50 | 106,869 |
| Estimated coverage** | 1043x |

\*Assembly input
\*\*Coverage based on Q155 chloroplast size

**Figure 1.**
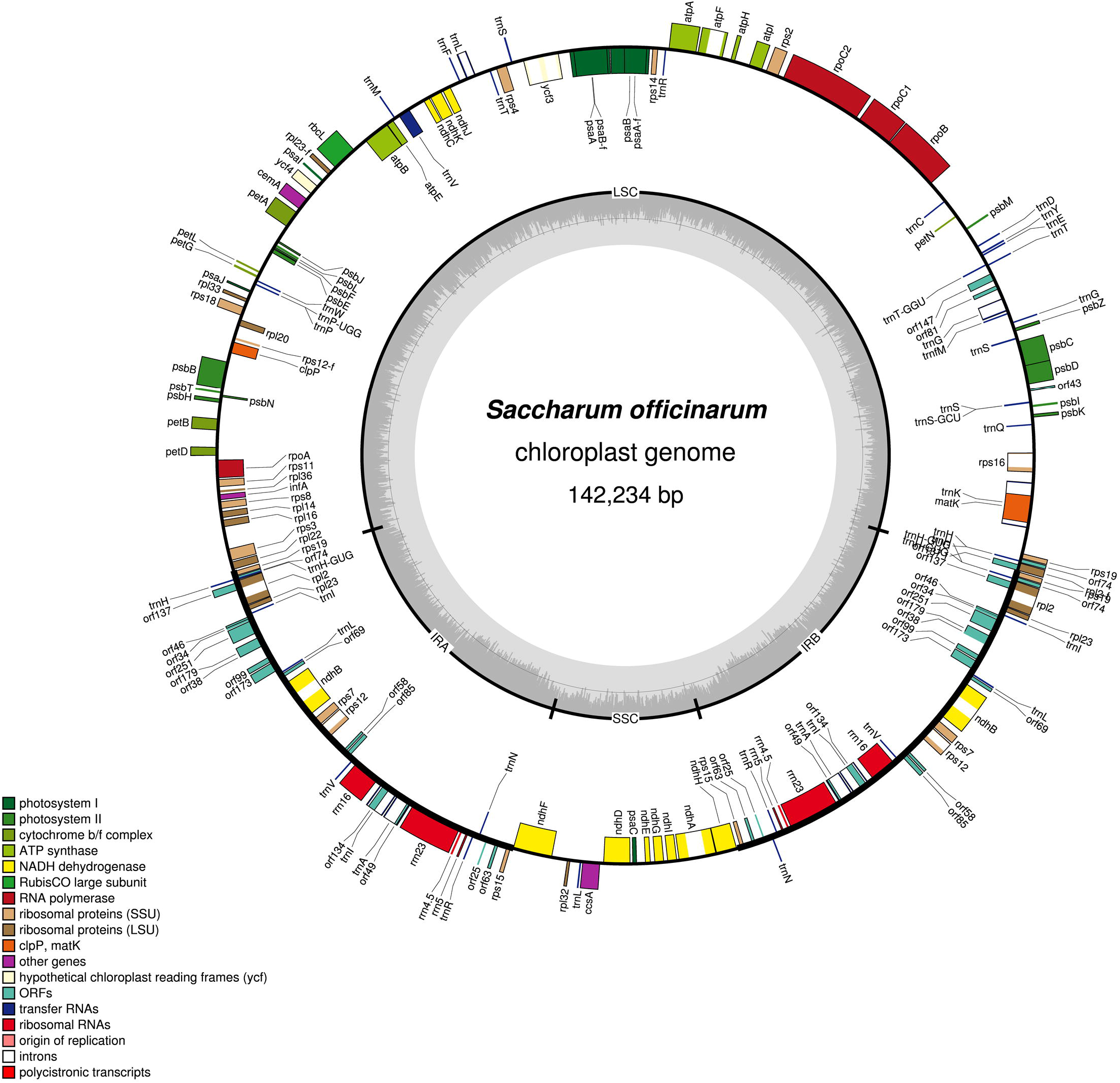
Graphic representation of *S. officinarum* chloroplast genome. Genes and tRNAs elements are identified as coloured boxes. The genes transcription direction is indicated by gray arrows. The locations of large and small single-copy regions and, the pair of inverted repeats (IRa and IRb) are shown in the inner circle. The darker gray color in the inner circle corresponds to the GC content, and the lighter gray color corresponds to the AT content.

The analysis of of the *S. officinarum* cpDNA revealed a notable difference in the LSC region of wild and cultivated sugarcanes. Chloroplasts of sugarcane cultivars such as *Saccharum* spp. Q155 (Hoang *et al*. 2015), *Saccharum* spp. NCo 310 (Asano *et al*. 2004), *Saccharum* spp. SP80-3280 (Calsa Júnior *et al*. 2004) and *Saccharum* spp. RB867515 (Vidigal *et al*. 2016) showed a loss of a duplicated fragment with 1,031 bp in the beginning of the LSC region. In comparison with those cultivars’ chloroplasts, *S. officinarum* has an insertion of 10 bp inside the *rp123-F* gene and two copies of *orf137, trnT, orf74* and *rps19* genes. Like the NCo310 chloroplast, *S. officinarum* chloroplast has an intron in the middle of the *rp12* gene. Based on these results, we propose the comparative analysis of organelle genomes as a very important tool for deciphering and understanding hybrid Saccharum lineages.

## ACKNOWLEDGEMENTS

We are grateful to Instituto Agronômico de Campinas for providing the wild species plant materials. We are thankful to FAPERJ (Fundação de Amparo à Pesquisa do Estado do Rio de Janeiro), CNPq (Conselho Nacional de Desenvolvimento Científico e Tecnológico) and CAPES (Coordenação de Aperfeiçoamento de Pessoal de Nível Superior) for the financial support.

